# Antimicrobial blue light-bathing therapy for wound infection control

**DOI:** 10.1101/2024.04.13.589323

**Authors:** Jie Hui, Wonjoon Moon, Pu-Ting Dong, Carolina dos Anjos, Laisa Negri, Hao Yan, Ying Wang, Joshua Tam, Tianhong Dai, R. Rox Anderson, Jeremy Goverman, Jeffrey Gelfand, Seok-Hyun Yun

## Abstract

The prevalence of antibiotic resistance and tolerance in wound infection management poses a serious and growing health threat, necessitating the exploration of alternative approaches. Antimicrobial blue light therapy offers an appealing, non-pharmacological solution. However, its practical application has been hindered by the requirement for high irradiance levels, which particularly raises safety concerns. Here, we introduce a light-bathing strategy that employs prolonged, continuous exposure to blue light at an irradiance range lower by more than an order of magnitude (5 mW/cm^2^). This method consistently applies bacteriostatic pressure, keeping wound bioburden low, all while minimizing photothermal risks. Leveraging tailor-made, wearable light-emitting patches, we conducted preclinical trials on rat models of wound infection, demonstrating its safety and efficacy for suppressing infections induced by methicillin-resistant *S. aureus* and multidrug-resistant *P. aeruginosa*. Our results pave a new way for the application of blue light therapy in wound care.

---

Wound infection is a major worldwide healthcare burden^1–3^. The requirement of orchestrated healing factors and the easy exposure to microbial contaminants render wounds, especially chronic wounds, susceptible to infection^3–6^. If the infection is not properly managed, it can lead to delayed healing, and serious or even life-threatening complications^3,7^. Both systemic antibiotics and topical antimicrobials are crucial for treating invasive infections. However, they encounter significant challenges in wound infection management, particularly associated with antimicrobial resistance, biofilm formation, and adverse effects. It is estimated that over 70% of bacteria responsible for wound infections are resistant to at least one common antibiotic^8,9^. This situation is further complicated by the formation of bacterial biofilms, which are able to display remarkably high antibiotic tolerance compared to their planktonic counterparts^10–12^. Current clinical guidelines advise the careful use of topical antimicrobials and systemic antibiotics and recommend limiting antiseptics to short-term use to avoid toxicity and hypersensitivity reactions^13–16^. These challenges highlight the urgent need for the development of innovative antimicrobial strategies.

Current antibiotic discovery has encountered significant obstacles evidently as no new class of antibiotics has been introduced into the clinic in the last four decades^17,18^. At the same time, antimicrobial resistance has become a paramount public health crisis^19–22^, contributing to 1.27 million deaths globally in 2019 and projected to 10 million annually by 2050^2^. The dwindling antibiotic discovery combined with the rising antimicrobial resistance have already propelled us into a post-antibiotic era. In response, the scientific community has investigated various alternatives, particularly those with novel antimicrobial mechanisms, minimized selective pressure, or reduced off-target issues. Techniques encompassing anti-virulence therapies, antimicrobial peptides, phage therapy, and antibodies have shown promise in early research^18,23,24^, yet their clinical adoption remains elusive. Among these non-antibiotic approaches, antimicrobial blue light therapy has received significant attention. This approach targets endogenous chromophores in bacteria, particularly porphyrin derivates, and produces bacteria-specific cytotoxic effects upon blue light illumination (400∼470 nm). Owing to its non-pharmacological nature, broad-spectrum efficacy, and no reported resistance, blue light therapy is considered a promising avenue^25–33^ for localized antimicrobial use, especially in wound care. Extensive laboratory and animal studies indicate its antimicrobial potency on many common wound pathogens^32–41^. Nevertheless, its clinical translation has faced substantial obstacles due to its limited *in vivo* bactericidal effectiveness and high photothermal risks due to high-irradiance/intensity illumination.

Current blue light therapy protocols aim to rapidly eliminate bacteria using a single or multiple doses, each lasting from 10 minutes to an hour, necessitating high optical intensities exceeding tens of milliwatts (mW) per cm^2^. Intensities between 50-100 mW/cm^2^ have shown potent bactericidal efficacy *in vitro* in a dose-dependent manner^29,33,34,36,38,39^. However, when advanced to *in vivo* testing on animal models, even 200 mW/cm^2^ — considered the Maximum Permissible Exposure (MPE) level given by the American National Standard Institute (ANSI) guidelines^42^ — at 300 J/cm^2^ levels results in only moderate (< 2-log_10_) reductions^32,41^. These high-intensity levels pose a serious risk of photothermal damage, more prominent under repeated dosing to prevent bacterial relapse^32,41,43^. Moreover, the demand for high irradiance places substantial practical constraints on light sources, especially for large wounds which requires high total power and proper thermal management for large-area illumination. This requirement can present an even bigger challenge for wearable development^44,45^.

Here, we introduce a promising solution to this bottleneck, challenging traditional approaches. We suggest that antimicrobial blue light therapy may not be best suited for quickly eliminating bacteria in infected wounds. Instead, its most effective use could be as a prolonged intervention during wound healing, applying continuous antimicrobial pressure on wound bed to maintain its bacterial load below a clinically acceptable level. This revised strategy, termed “light-bathing”, involves continuous or quasi-continuous exposure to blue light in a low-irradiance paradigm. In this paper, we present highly encouraging results from our *in vitro* and *in vivo* studies of this light-bathing approach, utilizing custom-built wearable light-emitting patches on rat models of wound infection. We discovered that prolonged exposure to 410-nm light at just 5 mW/cm^2^ can effectively prevent bacterial growth and keep the *in vivo* wound bioburden low for both gram-positive *S. aureus* and gram-negative *P. aeruginosa*, two most common antibiotic-resistant wound pathogens, over the entire 2-day treatment course. Its safety and effectiveness were rigorously confirmed through various methods, including clinical wound signs, immunohistochemical analysis, bioluminescence imaging, fluorescence *in situ* hybridization imaging, and endpoint colony-forming unit (CFU) enumeration from wound biopsies. Our findings highlight the potential of blue light-bathing as a new antimicrobial strategy in wound care.

## Results

### Minimum irradiance levels for bacterial growth inhibition

In conventional antimicrobial discovery and susceptibility test, the “Minimum Inhibitory Concentration” (MIC) is a widely used benchmark to characterize the lowest concentration of an antimicrobial agent that visibly inhibits the growth of a microorganism on agar plates or in broth medium^46^ (refer to Fig. 1a). Typically, an antibiotic’s effectiveness is gauged by maintaining a concentration at MIC and higher, with dosages adjusted as necessary. Drawing inspiration from this established metric, we introduced the concept of “Minimum Inhibitory Irradiance” (MII), which we define as the minimum irradiance of blue light required under continuous long-term illumination (e.g., overnight, or longer) to enable complete growth inhibition of a microorganism within an *in vitro* growth environment. To quantify MII, we conducted an *in vitro* assay involving the incubation of bacterial cultures on agar plates followed by concurrent overnight 37 °C incubation and continuous 410 nm light illumination at varying irradiances as illustrated in Fig. 1a. The growth of methicillin-resistant *S. aureus* (MRSA USA300), a prevalent wound pathogen noted for its tolerance to 410 nm light^29^, was fully inhibited after 24-hour exposure with light irradiances at 2 mW/cm^2^ (Fig. 1b), while irradiances below 1 mW/cm^2^ permitted limited bacterial growth. Post-treatment, additional 24-hour culture in darkness showed no relapse within the illuminated zones (Fig. S1). Therefore, the MII for MRSA (USA300) was determined to be 2 mW/cm^2^. By the same method, the MIIs for *P. aeruginosa* (ATCC 27853) and *E. coli* (ATCC 25922) were determined at 3 mW/cm^2^ and 2 mW/cm^2^, respectively (Fig. 1c). These MII values are more than an-order-of-magnitude lower than the typical irradiances employed in traditional blue light therapy (Fig. 1d). At such low irradiances, light- bathing — continuous illumination over extended periods, can be employed as a new regimen to ensure a sustained and sufficient antimicrobial pressure.

**Figure 1.**
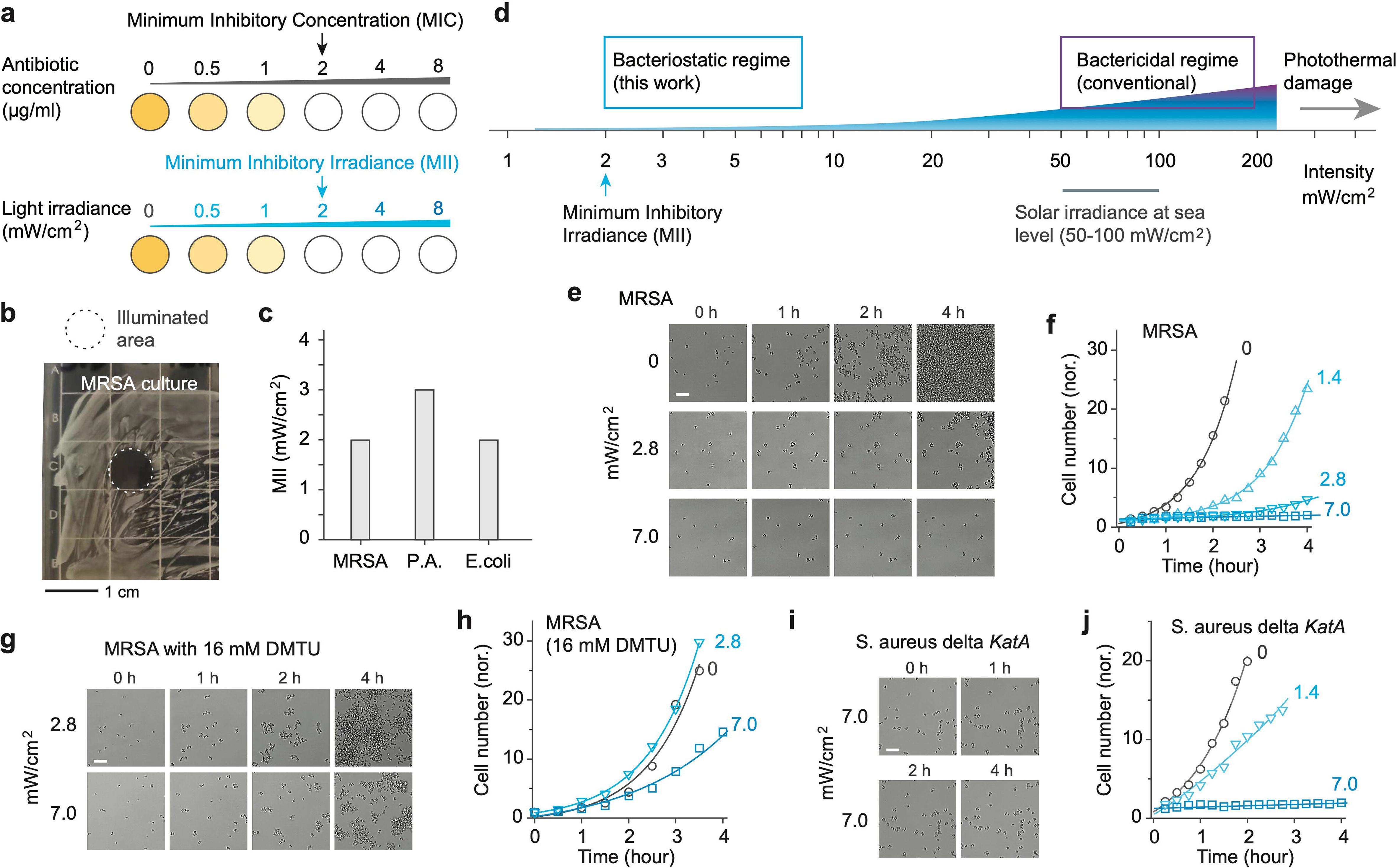
Minimum inhibitory irradiance (MII) and the proposed therapeutic window for bacteriostatic blue light therapy. (a) Illustration of MII, defined as the lowest irradiance that enables the complete bacterial growth inhibition under continuous 410 nm illumination, akin to the concept of Minimum Inhibitory Concentration (MIC) for antibiotics. (b) Image of a MRSA-streaked agar plate following overnight continuous exposure to 410-nm blue light at 2 mW/cm^2^, highlighting the treated area (dotted circle) through an 8-mm diameter aperture. (c) Measured MII levels for three prevalent wound pathogens: MRSA (USA300), *P. aeruginosa* (ATCC 27853), and *E. coli* (ATCC 25922). (d) Diagram illustrating the proposed bacteriostatic window employing significantly lower light intensities compared to traditional high-irradiance bactericidal approaches, which are comparable to solar irradiance at sea level under clear skies. (e) Time-lapse optical imaging of MRSA USA300 cells under continuous blue light exposure with varying irradiances. All cell groups underwent similar preparation and analysis. (f) Growth curves for MRSA, derived from the sequential images in (e). (g) Time-lapse imagery of MRSA USA300 cells treated with 16 mM DMTU to deplete reactive oxygen species (ROS). (h) Growth curves for the ROS-depleted MRSA. (i) Time-lapse imagery of catalase-deficient mutant *S. aureus* Δ*katA*. (j) Growth curves for *S. aureus* Δ*katA*. Each growth curve was analyzed using an exponential function to estimate cell doubling time. Scale bars represent 20 μm.

### Bacteriostatic effects of low-level light-bathing on *in vitro* culture

To delve deeper into the bacteriostatic effects of light-bathing, we observed the growth dynamics of various bacterial strains cultured in nutrient-rich brain-heart-infusion (BHI) media under continuous 410 nm illumination. In the absence of light, MRSA cells exhibited rapid proliferation (Video S1), with a cell doubling time of ∼29 minutes as deduced from the growth curve fitting (Figs. 1e and 1f). Continuous exposure to light at irradiances of 1.4 and 2.8 mW/cm^2^ resulted in a noticeable retardation of cell growth, extending cell doubling time to 52 and 128 minutes, respectively. At an irradiance of 7 mW/cm^2^, we observed a complete halt in cell proliferation. This level of irradiance surpasses the MII of 2 mW/cm^2^ identified for streaked agar plates, a variance attributed to both the rich nutrient and optical absorption in the medium (Fig. S2).

Similar to conventional high-irradiance blue light therapy^27,29,32^, the antimicrobial action at lower irradiances is presumed to stem from the generation of reactive oxygen species (ROS). To validate this hypothesis, we introduced N,N’-dimethylthiourea (DMTU), a ROS scavenger, into the culture medium at a concentration of 16 mM, and subsequently monitored the growth under varying light irradiances (Fig. 1g and Video S2). In the presence of DMTU, the difference in bacterial growth between the unilluminated control and the 2.8 mW/cm^2^ illuminated group was negligible. Notably, at 7 mW/cm^2^, bacterial proliferation was no longer inhibited effectively (Fig. 1h). This corroborates the ROS-mediated bacteriostatic mechanism of light-bathing.

Further exploration involved subjecting catalase-deficient mutant *S. aureus* (Δ*katA*) to light- bathing. Prior research has identified catalase as a critical molecular target for blue light, implicated in the disruption of H_2_O_2_ detoxification processes^31,32^. Contrary to our expectation, light-bathing exhibited comparable efficacy in both the catalase-mutant and methicillin-resistant *S. aureus* strains (Figs. 1i and 1j; Video S3). This observation suggests that there are other bacterial chromophores or mechanisms involved in the antimicrobial effect of light-bathing. While this aspect merits further investigation, the multi- molecular-target mechanism could implicate a reduced likelihood of resistance development.

### Animal model and light-delivering wound patch for *in vivo* testing

To extend our examination of the light-bathing strategy to live animals, a device capable of delivering light continuously to target sites is needed. We devised wearable patches equipped with blue light emitting diodes (LED) and engineered a tethered prototype optimized for use with freely moving rat models (Fig. 2a and Video S4). This device comprised a 10×10 LED array, covering a 22×22 mm^2^ area, emitting light at a spectrum of 409 ± 5.5 nm, with tunable output intensity (Fig. S3). To mitigate potential overheating of the device, we fitted it with a heat sink and a miniature cooling fan, enabling the device output to reach up to 200 mW/cm^2^ during biosafety assessment. However, we found that active cooling was not necessary at therapeutic irradiances below 20 mW/cm^2^, where without cooling, the LED’s output was stable, and device heating was negligible (Fig. 2b).

**Figure 2.**
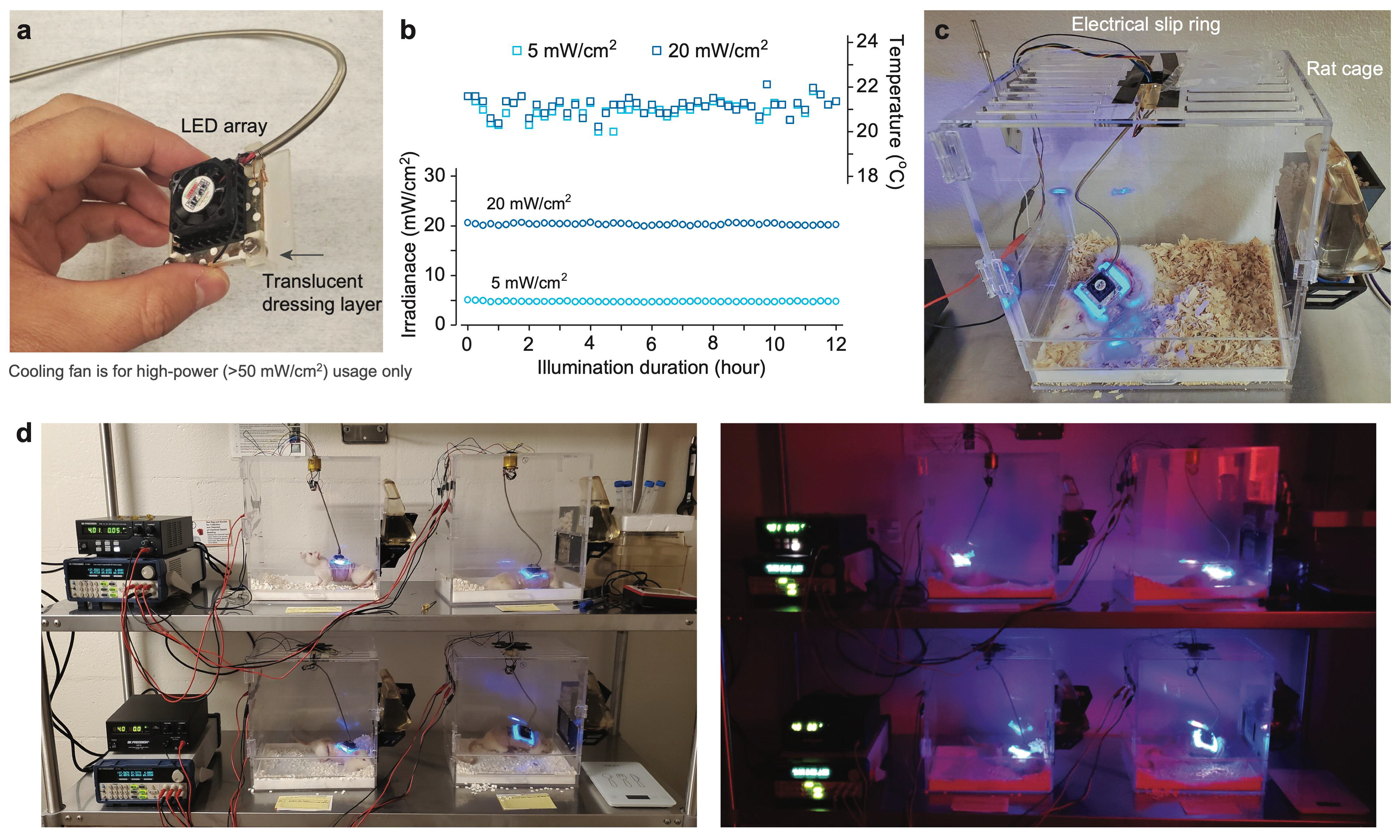
Design of antimicrobial photonic wound patch and prototype of tethered, wearable device for *in vivo* rat studies. (a) Image of the wearable light source featuring an LED array, cooling components for the LED, and a custom 3D-printed mounting frame. (b) Graph showing the stability of the LED’s output irradiance and temperature when operated without active cooling at 5 and 20 mW/cm^2^. (c) Photograph of the tethered, wearable device prototype fitted on a freely moving rat, illustrating the device in use with active blue-light illumination. (d) Overview of four complete sets of the device-cage system, including peripheral components, designed to enhance study throughput and validate reproducibility. Images capture rats under active light treatment, depicted with and without the presence of ambient lighting.

For animal trials, we selected the dorsal region of rats as the optimal site for both device placement and subsequent abrasion wound creation. Following the induction of wounds, we applied a standard transparent film drape before mounting the LED headpiece. The assembled wound patch was mounted onto intact dorsal skin, maintaining a distance of approximately 5 mm between the LED emitting facet and tissue (Fig. S4). To accommodate the unique requirements of our study, rat cages were customized to securely house the tethered wound patch device (Fig. 2c). The design and refinement of the cage setup were achieved through iterative testing and optimization. Particularly, a spring-loaded tether and an electrical slip ring were incorporated to reduce restrictions on rat mobility and prevent cable entanglement or chewing. We constructed four identical device-cage setups to enable simultaneous experimentation across multiple individual subjects (Fig. 2d).

### Phototoxicity evaluation on healthy and wounded skin

While MII sets the lower bound of the therapeutic window depicted in Fig. 1d, the risk of phototoxicity determines its upper bound. Our initial investigation focused on assessing the phototoxic effects of prolonged light exposure on healthy skin to establish the Maximum Permissible Irradiance (MPI), prioritizing photothermal safety. The schematic of our study protocol is presented in Fig. 3a. After preparatory steps of hair removal and skin cleansing, we equipped the rat’s dorsal skin with a light-transmitting (92%) drape film and an LED headpiece (Fig. 3b). We employed an irradiance de-escalation method. In an initial short-duration (6 hours) test, the animals were found to endure an irradiance of 30 mW/cm^2^ without showing significant signs of discomfort, edema, and erythema. Consequently, this irradiance level was selected as the benchmark for following study with further extended exposure. The rats underwent a continuous 2-day light exposure at varying irradiances, after which we monitored the skin’s response over the next three days, a duration deemed adequate to observe any delayed phototoxic effects^47^. This was followed by taking skin biopsies for histopathological examination. This 2-day exposure period matches the typical interval for changing wound dressings in clinical settings.

**Figure 3.**
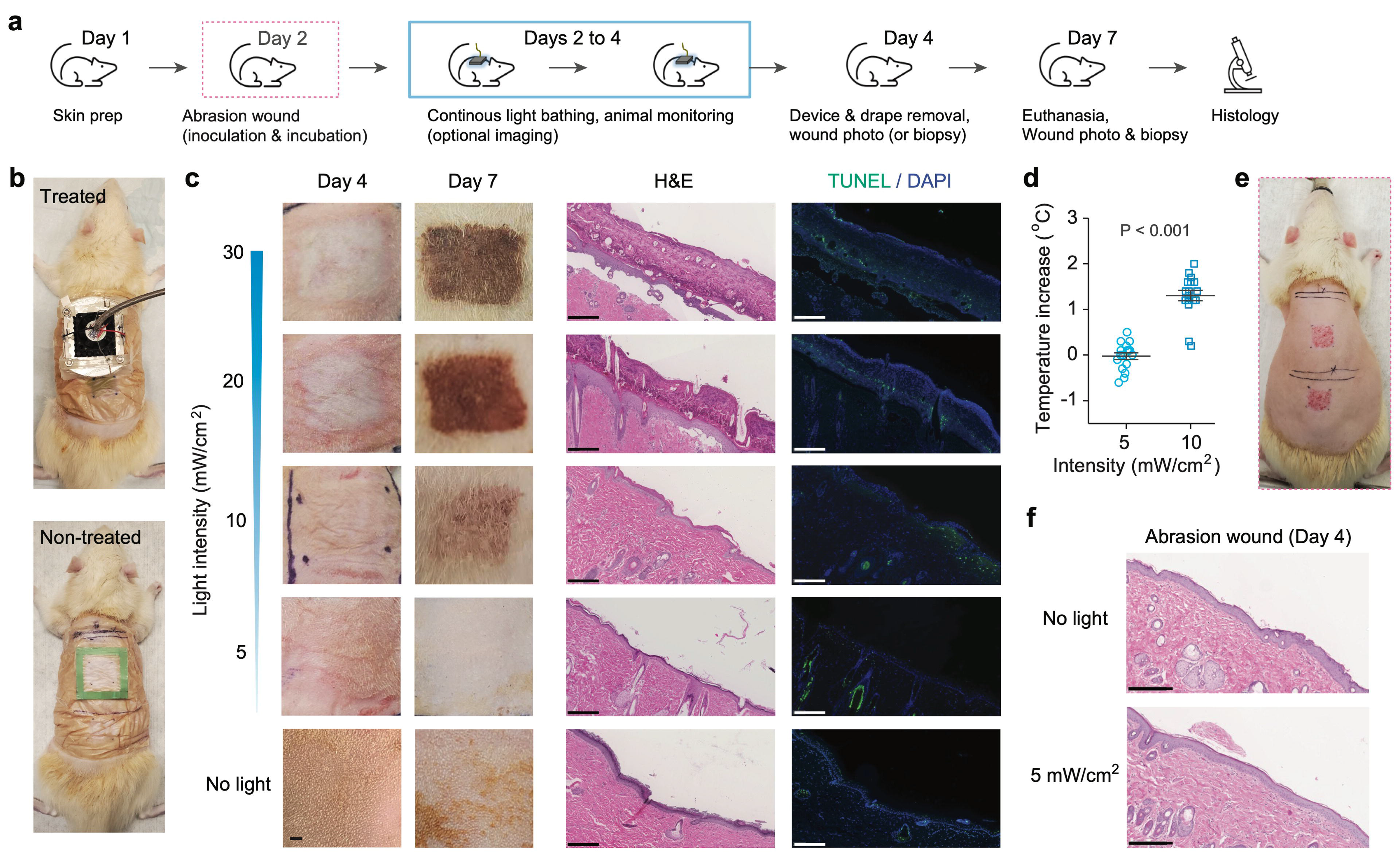
Evaluation of phototoxicity on *in vivo* rats. (a) Diagram detailing the phototoxicity study protocol conducted on healthy or wounded rat dorsal skin. (b) Comparative photographs depicting a rat equipped with the light-delivery device undergoing treatment (top) and a control rat not subjected to light exposure (bottom). (c) Bright-field *in vivo* images of the healthy skin regions captured on Days 4 and 7 after 2-day light treatment at different irradiances, accompanied by corresponding H&E-stained and TUNEL-stained histology images from biopsied samples collected on Day 7. (d) Graph illustrating the temperature increase measured at the dorsal skin surface during continuous illumination. (e) Images showing a rat with two freshly created abrasion wounds. (f) H&E-stained histological sections of dorsal abrasion wounds, comparing untreated versus 2-day light-treated at 5 mW/cm^2^. Scale bars indicate 250 μm.

Post-treatment, we observed no notable erythema or edema across all subjects, in line with Draize’s criteria used for toxicity test (Fig. 3c). However, animals exposed to higher irradiances of 20 and 30 mW/cm^2^ exhibited marked skin discoloration, indicative of phototoxicity affecting the epidermis. Three days after treatment, these discolored regions progressed to darkened, necrotic tissue layers (Fig. 3c), with this effect being less pronounced at 10 mW/cm^2^ and virtually absent at 5 mW/cm^2^. Further microscopic analysis through H&E and TUNEL staining elucidated the presence of 200-300 µm-thick necrotic layers and apoptotic cells at higher irradiance levels (Fig. 3c). Remarkably, samples exposed to 5 mW/cm^2^ showed no evidence of tissue necrosis or cellular apoptosis. Temperature measurements taken directly beneath the LED patch revealed no significant increase at 5 mW/cm^2^, whereas a modest rise of 1.3 °C was noted at 10 mW/cm^2^ (Fig. 3d). Based on these results, we set the MPI range for rat skin between 5 and 10 mW/cm^2^.

Further evaluations were conducted on abrasion-induced wounds on the rat’s dorsal skin (Fig. 3e), mimicking superficial injuries by removing the stratum corneum and upper epidermis (Fig. S5). In the absence of light exposure, these non-contaminated wounds naturally healed within 48 hours. Histological examination confirmed similar regeneration of the stratum corneum and epidermis in areas treated with light (Fig. 3f), indicating that 2-day light-bathing at 5 mW/cm² does not hinder the wound healing process. It is important to note that this irradiance level is 40 times lower than the ANSI MPE of 200 mW/cm², which has been frequently cited to justify higher irradiances in previous preclinical studies of blue light therapy. Our findings urge caution regarding the safety of using a few tens of milliwatts per cm² in blue wavelengths.

### Light-bathing therapy on MRSA-infected wound

To assess the bacteriostatic efficacy of light-bathing, we integrated contamination and infection phases into our abrasion wound model. We inoculated the wound surfaces, each measuring approximately 15 mm by 15 mm, with a high load (∼ 10^8^ CFUs) of MRSA USA300 immediately following the abrasion procedure, covered with a drape film, and allowed a 3-hour period for colonization and infection. Drawing on our MII and MPI values, we selected an irradiance of 5 mW/cm^2^ for all further evaluations. For observation of the dynamic infection evolution, we first employed a bioluminescent MRSA strain (MRSA USA300 *lux*^43^) and performed time-lapse bioluminescence imaging (BLI)^48^ to monitor the bacterial load over a 44-hour period, encompassing 41 hours of light exposure. Additionally, we infected another set of rats with non-luminescent antibiotic-resistant clinical isolate (MRSA USA300) following the same protocol and then conducted punch biopsies on wounds^49^ for comprehensive bioburden and wound condition analyses.

The BLI signal, proportional to bacterial load with a detection limit of 1.8×10^5^ CFU/cm^2^ for MRSA USA300 *lux*, indicated that the initial high inoculation load resulted in a robust BLI signal across the infected area (Figs. S6-8). In the control group, the bacterial load peaked around 12 hours post-inoculation, then gradually declined, likely due to natural host defense mechanisms, although scattered hot spots of activity persisted at 44 hours (Fig. 4a and Fig. S9). Notably, one control animal exhibited a pus-like area at 12 hours and an ongoing active infection at the 44-hour mark (middle in Fig. 4a). By contrast, the bacterial load observed in the light-treated group in general started to decrease almost immediately after light treatment initiated and continued over time, culminating in nearly complete eradication by the 44-hour endpoint (Fig. 4b and Fig. S9). As apparent in the dynamic curves of bacterial load quantified via BLI (Fig. 4c), treated wounds showed significantly lower bioburden levels across all time points, especially notable when compared to the first 20 hours of active infection in the control group. Visual assessment further confirmed the cleaner and healthier appearance of treated wounds compared to the inflamed and pus-producing control wounds (Fig. 4d). Endpoint CFU bioburden analysis on wounds infected with non-luminescent MRSA showcased a substantial reduction in bacterial load in treated groups, a 2-log_10_ reduction compared to both control groups and initial inoculation levels (Fig. 4e). Fluorescence in situ hybridization (FISH) using *S. aureus*-specific peptide nucleic acid probes (PNA-FISH) highlighted a drastically decreased bacterial presence, primarily confined to the surface of treated wounds (Fig. 4f and Fig. S10).

**Figure 4.**
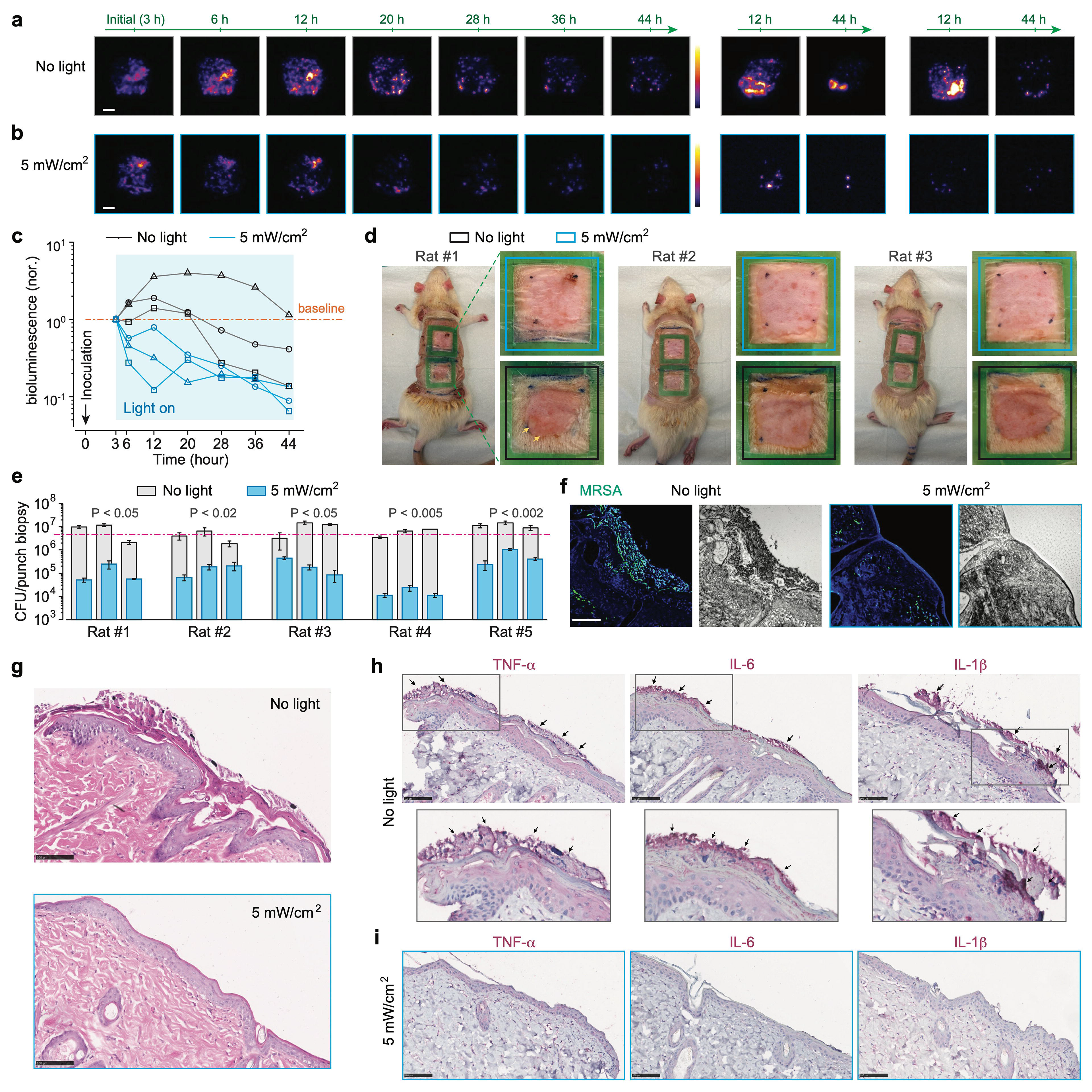
Evaluation of effectiveness on *in vivo* rat models of MRSA wound infection. (a-b) Time-lapse bioluminescence imaging shows the progression of MRSA USA300 *lux* infection in rat dorsal abrasion wounds, comparing untreated (a) to those receiving 2-day light therapy at 5 mW/cm^2^ (b). The three sets of images per group are acquired from three independent replicates. (c) Plots of relative integrated bioluminescence intensity of each wound, which is normalized to the initial value at 3 hours, illustrating the dynamic bacterial load over time. (d) Endpoint visual assessment of wound appearance. Yellow arrows indicate infection-induced pus. (e) Endpoint CFU enumeration of bacterial load within punch biopsy samples taken from each wound. Pink dashed line indicates the initial inoculation load. (f) Endpoint *S. aureus*-specific PNA-FISH imaging provides detailed visualization of bacterial presence in punch biopsy samples from each group. (g) Endpoint H&E-stained histological sections highlight tissue morphology. (h-i) Endpoint immunohistochemical staining of several key cytokines in samples from the untreated group (h) versus the light-treated group (i). Arrows indicate the extensive expression of corresponding cytokine markers in the untreated group. Scale bars, 5 mm for (a) and 100 μm in (f-i).

Morphological changes and inflammatory response evaluated through H&E and immunohistochemical staining for TNF-α, IL-6, and IL-1β cytokines of the biopsied samples further delineated the therapeutic effect. Control wounds exhibited typical signs of infection-induced inflammation, including necrotic tissue discharge and epidermal proliferation (Fig. 4g). In stark contrast, treated wounds displayed no such pathology, appearing virtually identical to normal, healthy skin. The absence of pro-inflammatory cytokines in treated samples underscored the therapy’s ability to not only control MRSA infection but also facilitate the healing process by mitigating inflammation (Figs. 4h and 4i). Through these analyses, light-bathing therapy has demonstrated profound success in managing MRSA-infected wounds.

### Light-bathing therapy on *P. aeruginosa*-infected wound

Turning our attention to *P. aeruginosa*, particularly known for its virulence and multidrug resistance, we utilized two different strains, a bioluminescent strain (PAO1 *lux*^28,50^) for time-lapse infection monitoring and a clinical strain (CDC AR Bank #0231) for endpoint analyses. PAO1 *lux* exhibits a *lux* expression more than 200 times higher than the MRSA USA300 *lux* strain, offering a minimum detection sensitivity of 7.4×10^2^ CFU/cm^2^. In untreated wounds (Fig. 5a and Fig. S11), the infection indicated by the measured bioburden escalated, peaking at 12 hours and subsequently leading to the extensive formation of wound exudate (pus), signs indicative of an aggressive, invasive infection. Remarkably, the application of light- bathing at 5 mW/cm^2^ not only curtailed this escalation but progressively reduced the bacterial load over time, as evidenced by the apparently diminishing BLI signal (Fig. 5b and Fig. S11), without any noticeable pus formation. This contrast highlights the antimicrobial effectiveness of our light therapy by significantly suppressing the bacterial population below the initial inoculation levels throughout the treatment duration (Fig. 5c). Visual inspections further differentiated the outcomes between treated and untreated groups. Non-treated wounds were marred by greenish, spreading exudate visible through the transparent film drape, whereas treated wounds maintained a much cleaner and healthier appearance (Fig. 5d).

**Figure 5.**
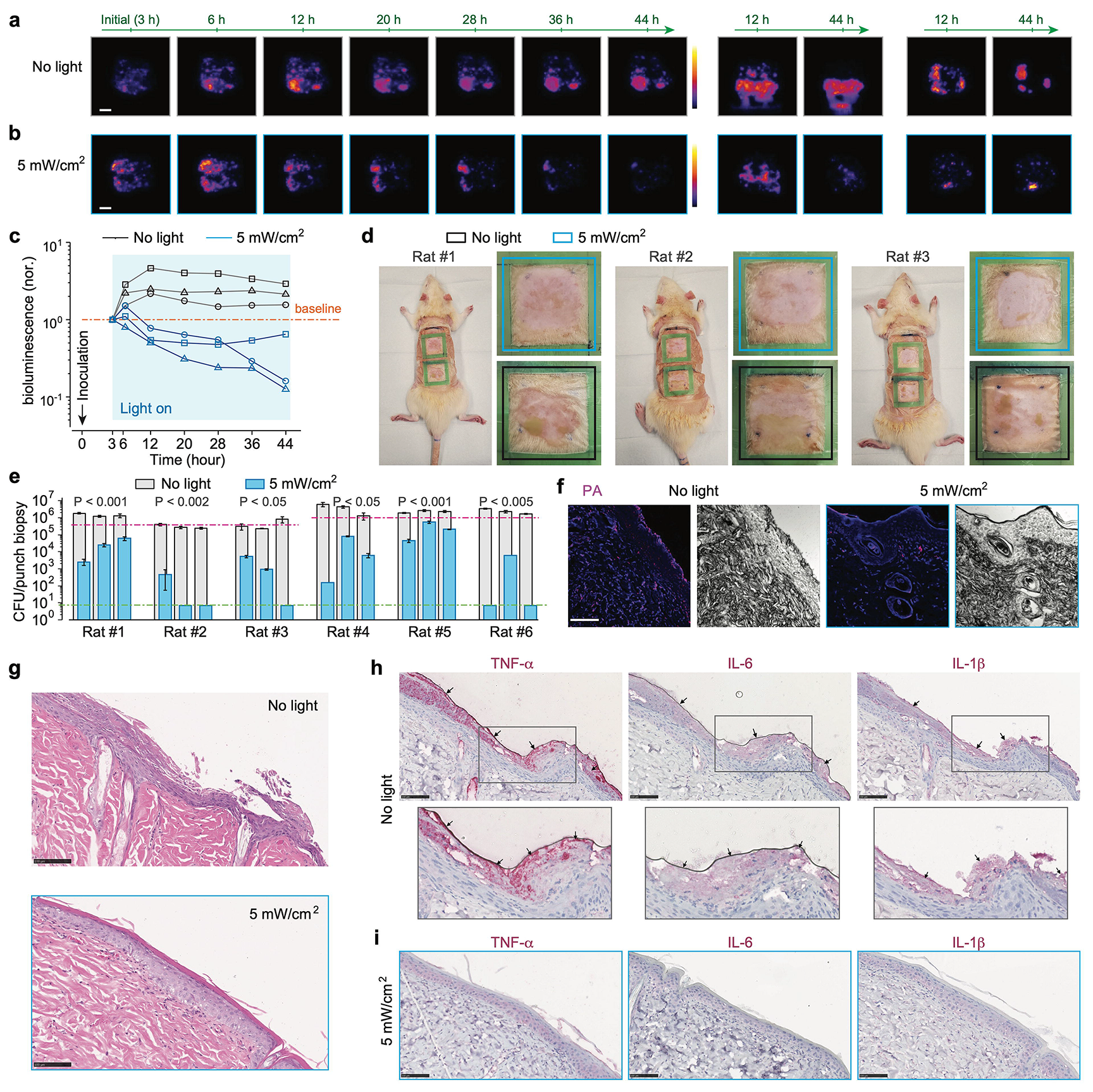
Evaluation of effectiveness on *in vivo* rat models of *P. aeruginosa* wound infection. (a-b) Time-lapse bioluminescence imaging tracks the progression of *P. aeruginosa* infection in rat dorsal abrasion wounds, showcasing untreated wounds (a) against those subjected to 2-day light therapy at 5 mW/cm^2^ (b). The three sets of images per group are acquired from three independent replicates. (c) Plots of relative integrated bioluminescence intensity of each wound, which is normalized to its initial value at 3 hours, illustrating changes in bacterial load over time. (d) Endpoint visual assessment of wound appearance. Extensive wound exudates are observed in the untreated wounds. (e) Endpoint CFU enumeration of bacterial load within punch biopsies randomly sampled from each wound. Pink and green dashed lines indicate the initial inoculation load and the detection limit, respectively. (f) Endpoint *P. aeruginosa*-specific PNA-FISH imaging for detailed visualization of bacterial distribution in punch biopsy samples from each group. (g) Endpoint H&E-stained histological sections. (h-i) Endpoint immunohistochemistry results of key cytokine markers in samples from the untreated groups (h) compared to the light-treated group (i). Arrows indicate the extensive expression of corresponding cytokine markers in the untreated group. Scale bars indicate 5 mm in (a) and 100 μm for (f-i).

Following the same protocol, we used a multidrug-resistant *P. aeruginosa* clinical isolate (CDC AR Bank #0231) and performed similar bioburden and wound condition analyses. Quantitative analysis through CFU enumeration of wound biopsies revealed a reduction exceeding 3-log_10_ CFU on average in the treated group compared to the untreated control (Fig. 5e), with several samples achieving ‘sterilization’ levels of bacterial reduction, as their CFUs were diminished beyond the detection limit. This result was corroborated by *P. aeruginosa*-specific PNA-FISH imaging, which showed a dramatic bacterial reduction, particularly in the superficial epidermal layers of treated wounds, in contrast to the dense bacterial presence in untreated wounds (Fig. 5f and Fig. S10). Histological evaluations provided further insights into the therapeutic impact of light-bathing. H&E staining of non-treated wounds displayed thick, infected lesions across the wound surfaces, whereas treated wounds exhibited normal skin morphology, free from infection-induced disruptions (Fig. 5g). The absence of inflammatory markers such as TNF-α, IL-6, and IL-1β in treated wounds (Fig. 5i), as opposed to their extensive expression in untreated wounds (Fig. 5h), confirmed the dual antimicrobial and anti-inflammatory efficacy of light-bathing therapy in managing *P. aeruginosa*-infected wounds.

## Discussion

Our research represents a substantial progress in antimicrobial blue light therapy by introducing a novel bathing regimen that diverges from the traditional protocols which utilize high irradiance levels (50-200 mW/cm^2^) and short treatment durations (10-60 min). The identification of a new therapeutic window for blue light therapy is crucial, highlighting its biosafety, effectiveness, and feasibility for clinical use. Contrary to conventional high-irradiance methods aiming at rapid and complete bacterial eradication, our approach sustains the necessary antimicrobial pressure to maintain bacterial loads below clinically acceptable levels (e.g., < 10^5^ CFUs/g) throughout the treatment.

A limitation of our current study is its focus on acute wounds at an early infection stage. Future research is warranted to assess the therapy’s effectiveness against more advanced infections, especially those involving biofilms. Nonetheless, the most practical and significant application of blue light therapy is on wounds prepared by clinical procedures that disrupt biofilms and remove necrotic tissues, optimizing wound bed for healing^13,51,52^. Such procedures are routine particularly in the management of chronic wounds and in preparing wound bed for skin grafting^53,54^. With biofilms disrupted and dispersed on wound surface, it creates a window to employ blue light therapy for wound infection control. While our study demonstrated the feasibility of continuous light exposure for 2 days, extending the treatment from several days up to several weeks could be achievable without significant phototoxicity. Investigating the impact of various intermittent illumination patterns, such as 16 hours on followed by 8 hours off, could also prove beneficial to accommodate the diverse needs and circumstances of patients.

The utilization of low irradiance levels, just several milliwatts per cm^2^, significantly aids the development of photonic wound patches. Negative pressure wound therapy (NPWT)^51,52,55,56^ is a common treatment for chronic wounds and extensive traumas prone to infection. Integrating light-bathing with NPWT devices could offer sustained infection control and accelerate wound healing. While gallium nitride-based blue LED technology is suitable for many cases, exploring flexible organic LEDs, fiber-optic patches, and wireless-powered light sources^57–59^ opens up innovative possibilities to accommodate various wound shapes, treatment lengths, and patient mobility needs.

## Methods

### Bacterial strains and chemicals

This study utilized bacterial strains including MRSA (USA300), MRSA (USA300 LAC::*lux*), *S. aureus* Δ*katA* (sourced from the George Liu Lab at UCSD), *P. aeruginosa* (PAO1 LAC::*lux*), *P. aeruginosa* (CDC AR Bank #0231), *P. aeruginosa* (ATCC 27853), and *E. coli* (ATCC 25922). Chemicals employed were N,N’-dimethylthiourea (DMTU) (D188700, Sigma-Aldrich), brain heart infusion (BD Difco 237500, Fisher Scientific), mannitol salt agar (1054040500, Sigm-Aldrich), cetrimide agar (OXCM0579B, Thermo Scientific), poly-D-lysine hydrobromide (P0899, Sigma), M9 medium (A1374401, Thermo Fisher), and Mueller-Hinton agar (R454082, Thermo Fisher).

### Bacterial culture condition

Bacterial strains were preserved in brain heart infusion (BHI) media enriched with 20% (v/v) glycerol at - 80°C. For experimental use, frozen stocks were thawed and streaked on BHI agar plates, then incubated overnight at 37°C to facilitate colony development for future use. Stationary-phase bacterial inoculum was prepared by transferring colonies from previously prepared streaked agar plates into sterile BHI media and incubating them in an orbital shaker set at 100 rpm overnight at 37°C. Prior to experimentation, bacterial cells were pelleted by centrifugation, washed twice with phosphate-buffered saline (PBS), and resuspended in 1X PBS.

### Light sources

For illumination of *in vitro* samples, as well as for the photothermal heating measurements on rat dorsal skin *in vivo*, we utilized an LED-based illumination system. This system comprised a single-element LED (M405L4, Thorlabs), equipped with an adjustable collimation adapter (SM2F32-A, Thorlabs) for beam focusing, powered by a dedicated LED driver (LEDD1B, Thorlabs) connected to a power supply unit (KPS201, Thorlabs). The LED emitted light centered at 409 nm within a 9 nm bandwidth, capable of producing a maximum output power of 1 W. The output power was variable and set to operate in continuous mode, with the beam size tailored via the collimation adapter for precise application. For *in vivo* rat studies, we used a 10×10 LED array (1DGL-JC-100W-405, Chanzon) with a center wavelength of 409 nm, an 11 nm bandwidth, and a peak output capacity of 100 W. This array covered a 22×22 mm^2^ area and provided an emission angle between 120-140 degrees, ensuring broad and even coverage. The irradiances and output powers of these lighting systems were measured by using a power meter (S121C and PA400, Thorlabs). Additionally, the emission spectra were evaluated with a spectrometer (CCS200, Thorlabs).

### *In vitro* bacteriostatic therapy

In our study of bacteriostatic effects of blue light, we conducted experiments on both agar plates and in broth medium. For the study on agar plates, stationary-phase bacterial inocula were uniformly streaked onto agar plates (e.g., Mueller-Hinton agar plates). These plates, covered with transparent lids, were then exposed to blue light from an LED device within a 37°C incubator. The illumination targeted a 1-cm diameter area on the plates and was maintained overnight (>10 hours) at varying irradiances. Post-illumination, the formation of bacterial colonies in both illuminated and non-illuminated zones was assessed visually. For the study in broth medium, stationary-phase bacterial inocula were introduced into diluted BHI medium (diluted 5× to minimize optical attenuation, refer to Fig. S2) and 30 μL of this bacterial suspension was allocated to each well of a 96-well plate. The plate was then placed in a 37°C incubation chamber on the sample stage of a laser-scanning confocal microscope (FV3000, Olympus), where blue light from multiple single-element LED sources was directed on to the wells through an auxiliary port of the microscope.

### Temperature measurements

To assess the thermal impact of blue light exposure, we conducted temperature measurement on *in vivo* rat skin exposed to air using an infrared thermometer (DT8011H, Thermco Products). Rat dorsal skin hair was removed a day prior to the experiment. Under anesthesia, two distinct dorsal skin areas (each 2×2 cm^2^) were illuminated with two separate LED beams set at 5 and 10 mW/cm^2^, respectively, while adjacent, non-illuminated skin areas served as controls. Additionally, the temperature of the LED arrays without active cooling components during continuous illumination was monitored using the same infrared thermometer.

### Confocal imaging

For our time-lapse confocal imaging studies, we utilized a setup integrating a single-element LED with a laser scanning confocal microscope (FV3000, Olympus). Bacterial suspension allocated into a 96-well plate was positioned within an incubation chamber on the microscope. The LED’s collimated beam was precisely aimed at targeted wells, with special attention to calibrating the irradiance reaching the bacterial suspension in the wells considering the optical attenuation caused by both the incubator window and the plate lid. Time-lapse images were captured using an UPlanFLN 20× objective (Olympus, NA 0.5) and subsequently processed and analyzed with ImageJ software.

### Animals

All animal experiments conducted in this study received approval from the Institutional Animal Care and Use Committee (IACUC) of Massachusetts General Hospital, ensuring compliance with the National Institutes of Health guidelines (Approval Number: 2022N000039). Male Sprague Dawley rats (CD® Sprague Dawley IGS, 11 weeks old) were sourced from Charles River Laboratories (Massachusetts, USA).

### Wearable photonic wound patch devices

For our *in vivo* studies with rats, we designed tethered wearable wound patches incorporating a 10×10 LED array (1DGL-JC-100W-405, Chanzon). To manage heat, a heat sink (345-1103-ND, Digi-Key) was affixed directly to the LED array’s backing metal plate. Additionally, a compact cooling fan (1570-1113- ND, Digi-Key) was secured atop the heat sink using super glue for enhanced thermal management. Electrical wiring connected to the LED and fan was fully encased within a metal spring (PS115H, Instech Laboratories) for protection and flexibility. The proximal end of the spring was secured to the LED head piece with super glue and metal wires, with its distal end connecting to an electrical slip ring (ZC- MOFLON-Slipring, Taida), facilitating rotation without wire tangling. This slip ring, featuring a 12.7 mm inner hole and a 33 mm outer diameter, accommodated up to 6 independent wires, ensuring reliable connection to a power supply (22-9130C-ND, Digi-Key). Customized rat cages were developed to accommodate the rotary slip rings, the wearable light devices, and the rat’s feeding, with the slip ring’s stationary side mounted on the rat cage. A four-pin quick connector streamlined the attachment and detachment process of the wound patch system from the cage. For animal studies, abrasion wounds were initially covered with a transparent film drape (V.A.C. Drape, 3M) to maintain sterility. A customized plastic splint was utilized to stabilize the wound area beneath the drape, ensuring optimal alignment of the LED array with the wound site. The LED device was mounted on rat by using a custom 3D-printed frame made from high-temperature V2 resin (Formlabs), which maintained a 5-mm gap between the LED output facet and the tissue to avoid direct contact. This frame was securely fastened to the rat dorsum, ensuring stable positioning throughout the study.

### *In vivo* phototoxicity study on healthy skin

Phototoxicity of our blue light-bathing therapy was first assessed on healthy dorsal skin of rats. On the first day, under isoflurane anesthesia, rats had their dorsal and abdominal hair fur shaved, followed by the application of hair removal cream for complete hair removal. Subsequently, the dorsal skin was cleaned with 4% chlorhexidine gluconate solution before mounting a transparent film drape (V.A.C. Drape, 3M) and the LED device over the shaved area. Starting on the second day, the LED device was activated to deliver varying irradiances in a de-escalation sequence (from 30 to 20, 10, 5, and 0 mW/cm^2^ at the skin surface) over periods of continuous illumination ranging from 36 to 48 hours. Following a 6-hour period at 30 mW/cm^2^ with one rat showing good tolerance, we then extended our full-course illumination on this rat and followed by another rat at 20 mW/cm^2^. Subsequently, two rats were subjected to 10 mW/cm^2^, and five rats each were assigned to groups treated with 5 mW/cm^2^ and as a no-light control, respectively. After two-day exposure (at Day 4), the rats were re-anesthetized for device removal and initial wound condition documentation through photography. The animals were then returned to their cages for skin reaction observation over the next three days, culminating on Day 7. Final photographs were taken before the rats were euthanized for tissue biopsy collection. These samples were subjected to H&E and TUNEL staining for comprehensive histopathological analysis of the skin’s response to the light exposure.

### *In vivo* study on abrasion wounds

The impact of phototoxicity and therapeutic efficacy of the light-bathing therapy was assessed on rat dorsal skin subjected to abrasion wounds with or without induction of bacterial infection. Following the initial hair removal procedure identical to the healthy skin phototoxicity study, on the subsequent day, rats were anesthetized with isoflurane and received buprenorphine intraperitoneally for analgesia. Using a scalpel, we meticulously created two similar abrasion wounds (15×15 mm^2^ each), maintaining a separation of ∼20 mm along dorsal spine, by repeatedly scratching the skin to affect the stratum corneum and upper layer of the epidermis without damaging the dermis. Minimal to no bleeding was observed post-wounding. Custom plastic splints were applied around each wound to ensure a flat surface for consistent healing and light exposure. Each wound was then covered with a film drape that wrapped around the rat’s body, with an LED device positioned over the upper wound for 5 mW/cm^2^ light-bathing treatment. The placement near the head discouraged interference with the device’s wiring. The lower wound served as a control, receiving identical preparation without light-bathing treatment. After about 48 hours, the animals were euthanized, and the devices and drape were removed for follow-up analysis. This phase of the study utilized two rats, yielding four wound sites for examination.

For the investigation into the effects of light-bathing therapy on wounds with acute bacterial infection, overnight cultures of MRSA USA300 *lux*, MRSA USA300, *P. aeruginosa* PAO1 *lux,* or *P. aeruginosa* CDC AR Bank #0231 were prepared, washed in PBS, and adjusted to an optical density conducive to a bacterial inoculation load of approximately 10^8^ CFUs. The 20 μL aliquot of the bacterial suspension was spread evenly across each wound surface. After allowing the inoculum to dry, wounds were sealed with film drapes, and the LED device was affixed for subsequent light treatment. Rats were then housed in customized cages to allow for acute infection development over three hours. Following this infection period, light therapy commenced. Three rats for each bioluminescent strain and seven rats for each non-bioluminescent clinical strain were used (1 rat in *P. aeruginosa* CDC AR Bank #0231- infected group and 2 rats in MRSA USA300-infected group were dropped out due to device damage that occurred during light treatment), and three to five 3-mm punch biopsies were randomly sampled from each wound for subsequent tissue CFU enumeration, PNA-FISH imaging, or histopathological examination to assess the effects of the light treatment on wound healing and infection control.

### Bioluminescence imaging

*In vivo* BLI was executed using the IVIS Lumina In Vivo Imaging System (PerkinElmer). Initial tests on *in vitro* agar plates established the BLI signal’s linearity and minimum detection sensitivity for the bioluminescent strains. Considering the bioluminescence intensity variance between MRSA USA300 *lux* and *P. aeruginosa* PAO1 *lux*, optimal integration times were set at 240 and 24 seconds, respectively, for optimal signal-to-noise ratio without signal saturation. Bioluminescent strains were used to prepare rat models of abrasion wound infection, with three rats per group for treatment and control. BLI was conducted at various intervals up to 44 hours. For BLI imaging, rats were anesthetized and then positioned in the IVIS system equipped with anesthesia. The imaging process, including device removal and reattachment, was efficiently completed within 10 minutes. After imaging at the final time point, rats were euthanized, and wound conditions documented. BLI data were processed and analyzed with ImageJ software, providing critical insights into the dynamic bacterial presence within the wounds.

### CFU enumeration assay

To assess the viable bacterial load within each wound, three 3-mm punch biopsies were randomly sampled from each wound followed by a tissue CFU enumeration assay. The biopsy samples were collected into tissue lysing tubes (Lysing Matrix D, 2 mL tube, MP Biomedical) containing 800 μL of PBS. They were then lysed by using a high-speed tissue homogenizer (FastPrep-24 Classic Instrument, MP Biomedical) for thorough tissue disruption. Subsequently, 200 μL of the lysate was transferred to a 96-well plate for 10-fold serial dilutions to quantify CFUs for each sample. Mannitol salt agar (Sigma-Aldrich) and cetrimide agar (Thermo Fisher Scientific) plates were utilized for selective isolation and growth of *S. aureus* and *P. aeruginosa*, respectively. Plates were incubated at 37°C until visible colonies formed, allowing for CFU enumeration. For enhanced detection sensitivity, additional 100 µL of homogenate underwent glass bead plating across the entire surface of the agar plate, amplifying the detection sensitivity by 25-hold. Each CFU measurement included three technical replicates. The initial bacterial inoculation load in each rat experiment was similarly quantified. These measurements provided a comprehensive analysis of bacterial load both pre- and post-treatment.

### PNA-FISH imaging

Following protocols from prior studies^60,61^, we used Alexa488-tagged, 16S rRNA PNA probe to specifically target *S. aureus* and Cy5-tagged, 16S rRNA PNA probe to specifically target *P. aeruginosa*, developed in a peptide nucleic acid (PNA) format by PNA Bio Inc. Probe validation test was conducted first. Cultured MRSA USA300 or *P. aeruginosa* CDC AR Bank #0231 were fixed in 10% formalin for one hour and subsequently stored in 50% (v/v) ethanol at -20°C. Fixed cells were pelleted, rinsed, and resuspended in a hybridization solution containing dextran sulphate (10% wt/vol), NaCl (10 mM), formamide (30% v/v), sodium pyrophosphate (0.1% wt/vol), polyvinylpyrrolidone (0.2% wt/vol), Ficol (0.2% wt/vol), disodium EDTA (5 mM), Triton X-100 (0.1% vol/vol), Tris-HCl (50 mM, pH 7.5), and the respective PNA probe (200 nM). This mixture was incubated at 57°C for two hours for hybridization. Afterward, cells were centrifuged at 17,000 g for 6 minutes, resuspended in 500 µL of wash solution containing Tris Base (5 mM), NaCl (15 mM), and Triton X (1% vol/vol, pH 10), incubated at 57°C for 90 minutes, and then pelleted and resuspended in 1 mL sterile water. The prepared cells were placed on a glass slide, air-dried, and covered with ProLong Gold antifade mounting medium (Thermo Fisher Scientific, P36930) between the glass substrate and cover glass (VWR international, 48368-040). Imaging was performed with a Zeiss LSM780 confocal laser scanning microscope using a 63× oil immersion objective (NA 1.4) to capture Alexa488 and Cy5 fluorescence in a sequential-scanning mode with a field of view of 135 μm × 135 μm. With the successful probe validation, PNA-FISH hybridization for rat wound sections was further performed following a similar protocol but with hybridization conducted on UltraFrost adhesion slides (VWR international, 50-301-70). Paraffin-embedded punch biopsy sections from control and 5mW/cm^2^-treated wounds were placed on top of the UltraFrost adhesion slides followed by hybridization inside a temperature-controlled oven (57°C for 120 minutes). Post-hybridization with the hybridization buffer removed, wound sections were washed for 90 min at 57°C, air-dried, and further covered with ProLong Gold antifade mounting medium between the glass substrate and cover glass. Imaging was performed by utilizing the confocal microscope (Zeiss LSM780) with a UPlanFLN 20× objective (Olympus, NA 0.8), with a field of view of 425 μm × 425 μm, to visualize the fluorescently tagged bacterial cells within the wound sections.

### Histopathology and immunohistopathology

Histological and immunohistochemical analysis, including H&E, TUNEL, and immunohistochemistry, were conducted on punch biopsy samples for this study. Immediately after collection, these samples were fixed in 4% paraformaldehyde to preserve tissue architecture and cellular details. Following adequate fixation, the specimens were embedded in paraffin wax, and thin sections of 5 μm thickness were prepared and mounted on cover glasses for staining. The TUNEL staining, utilizing DeadEnd™ Fluorometric TUNEL System from Promega, was employed to identify apoptotic cells within the tissue by highlighting fragmented DNA. For the immunohistochemical staining, we targeted three pro-inflammatory cytokines: TNF-α, IL-6, and IL-1β. This was achieved using specific rabbit polyclonal antibodies: anti-TNF-α (ab6671, Abcam), anti-IL-6 (ab229381, Abcam), and anti-IL-1β (ab9722, Abcam). Following primary antibody incubation, sections were treated with a compatible secondary antibody (RALP525, Biocare Medical) and visualized using Warp Red Chromogen kit (WR806, Biocare Medical) to detect the antigens of interest. Finally, all stained slides, including those processed for H&E and TUNEL, were digitized using a NanoZoomer slide scanner (Hamamatsu) for detailed image analysis and documentation of histopathological changes.

### Statistical analysis

Data in this study are presented as mean ± standard deviation. Comparisons between two datasets were made using an unpaired *t*-test. A P-value < 0.05 was deemed to indicate a statistically significant difference, while P < 0.01 and P < 0.001 were considered indicative of even more significant differences. The notation ‘ns’ denotes a lack of significant difference.

### Data availability

The raw data underlying the graphs presented in this paper are accessible and will be provided upon a reasonable request to the corresponding author(s).

## Acknowledgements

This work is funded by the Air Force Office of Scientific Research via MMPP: FA9550-20-1-0063. R.R.A. was partially supported by the Lancer Endowed Chair in Dermatology. The authors thank Dr. Jenny Zhao with the Wellman Center Photopathology Core for histological analysis of biopsy samples, Dr. Sydney Cash lab for sharing rat cages, Sandeep Korupolu and William Farinelli for help in the design of the LED array devices, MGH Vaccine & Immunotherapy Center for offering a lab facility for *in vitro* experiments, and Dr. Haytham Kaafarani at the MGH Wound Center for providing clinical insights.

## Author contributions

J.H. and S.H.Y. designed the project. J.H. handled *in vitro* experiments, LED device development, and animal protocol creation. J.H. and W.M. performed rat studies. W.M. interpreted histology. P.T.D. performed FISH imaging. C.D.A. and H.Y. assisted in animal studies. L.N. helped *in vitro* experiments. Y.W. and J.T. contributed to animal protocols. T.D. contributed lab resources. R.R.A., J.T., J.G., and S.H.Y. secured funding. J.H., W.M. and S.H.Y. performed data analysis, prepared figures, and wrote the manuscript with input from all co-authors.

## Competing interests

The authors declare no conflict of interest.

## Notes

### Competing Interest Statement

The authors have declared no competing interest.

