## Supplementary Information for "Antimicrobial blue light-bathing therapy for wound infection control"

### Supplementary figures with legends

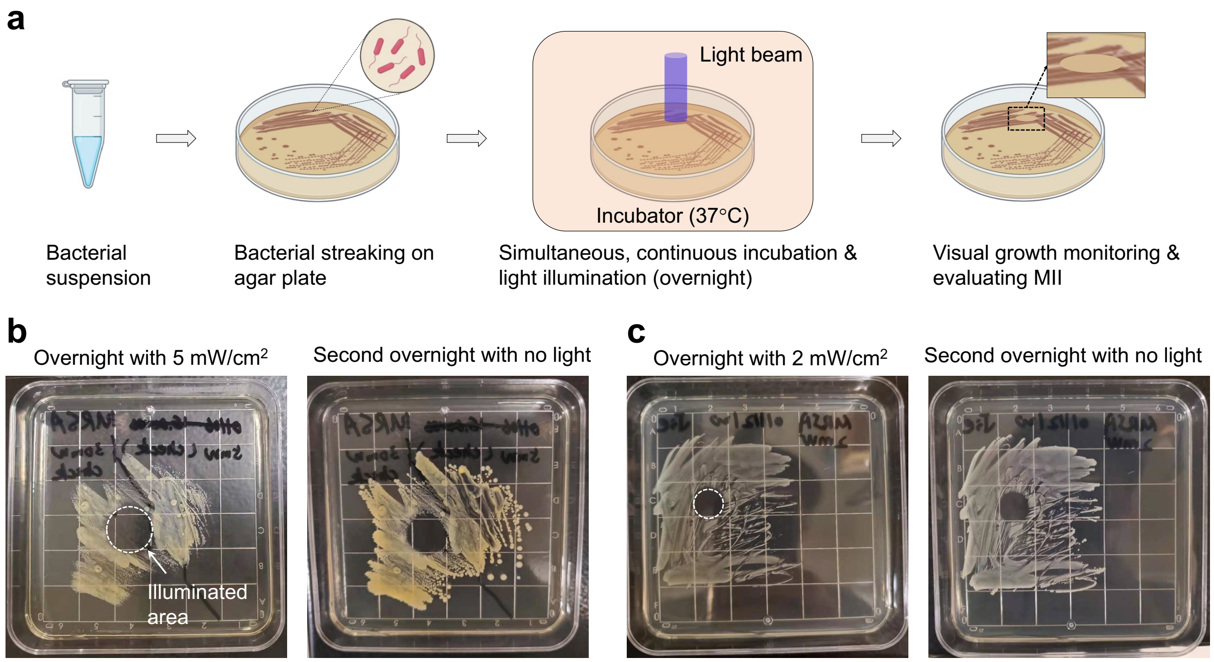


**Figure S1.** (a) Schematic of the protocol for measuring minimum inhibitory irradiance (MII) using *in vitro* streaked agar plates. (b) Bacterial colony formation on an agar plate after overnight incubation under 5 $mW/cm^2$ light (left), followed by another overnight incubation without light illumination (right). (c) Another agar plate showing bacterial colony formation after overnight incubation under 2 $mW/cm^2$ light (left), followed by another overnight incubation without light illumination (right). Plate grid size is 1 cm. The bacteria in the illuminated areas (dashed circles) are inhibited or eradicated while bacteria outside the illumination zones continue to proliferate.

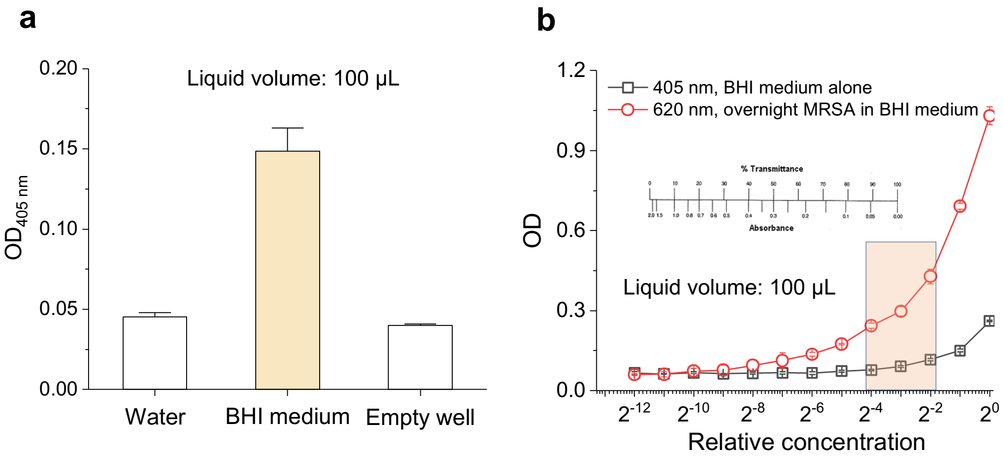


**Figure S2.** (a) Optical density (OD) at 405 nm measured from an empty well, 100 µL water in a well, 100 µL brain heart infusion (BHI) medium in a 96-well plate. (b) OD measurement at 405 nm of pure BHI media with different dilution factors, along with OD at 620 nm of overnight-incubated MRSA USA300-containing BHI media.

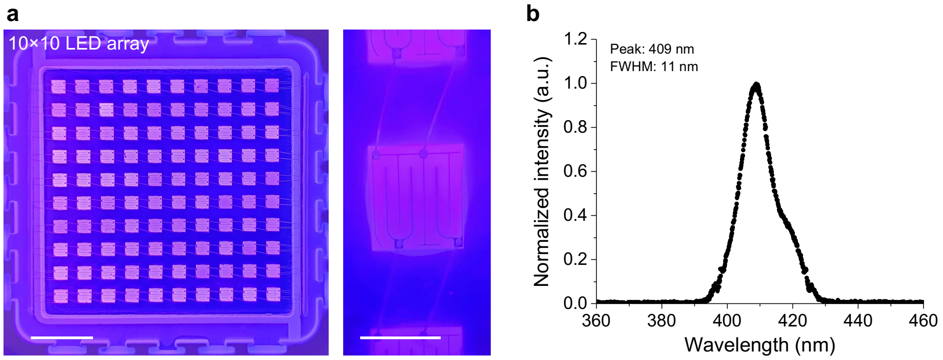


**Figure S3.** (a) Pictures of the LED array under active emission (1DGL-JC-100W-405, Chanzon). Scale bar, 4 mm (1 mm for the zoom-in image). (b) Emission spectrum of the LED array.

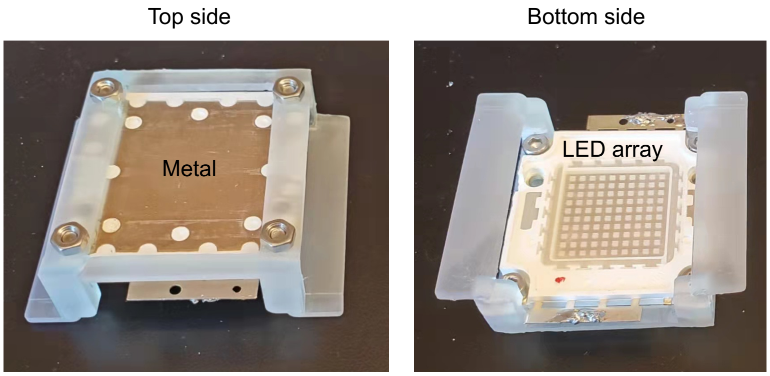


**Figure S4.** Design and pictures of the LED device mounting frame. The mounting frame was 3D printed using a high-temperature V2 resin and a 3D printer (both from Formlabs). Its height designed to keep a ~5 mm distance between the LED emission facet and skin/wound surface. Its positioning tab was used to firmly attach the mounting frame along with the LED device onto rat dorsum using stripes of adhesive film drape.

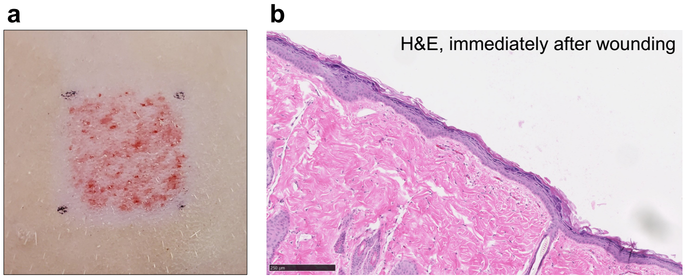


**Figure S5.** (a) Photo of a freshly induced abrasion wound on rat dorsal skin with an area of 15×15 mm$^2$. (b) Representative H&E histology image of the abrasion wound. Scale bar, 250 µm.

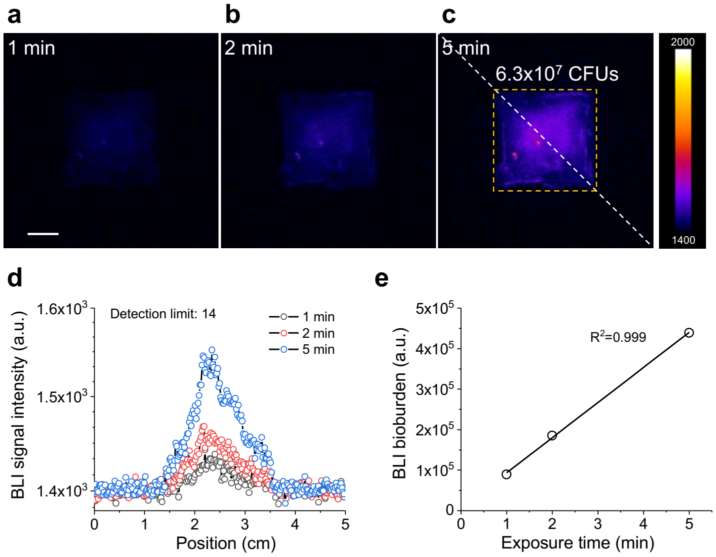


**Figure S6.** Measurement to show the linear relationship between bioluminescence (BLI) signal and signal integration time for MRSA USA300 *lux* cells. (a-c) Images of MRSA USA300 *lux* cells inoculated on an agar plate taken at different integration times. The inoculation load was 6.3×10$^7$ CFUs over an area of 15×15 mm$^2$ (yellow dashed frame in (c)) simulating the bacterial inoculation on a rat wound. Scale bar, 5 mm. (d) BLI signal plot for (a-c) along a diagonal axis (white dashed line in (c)). Its detection limit was 14 CFUs under the imaging setting. (e) The integrated BLI signal intensity or total BLI bioburden for (a-c) over the integration time.

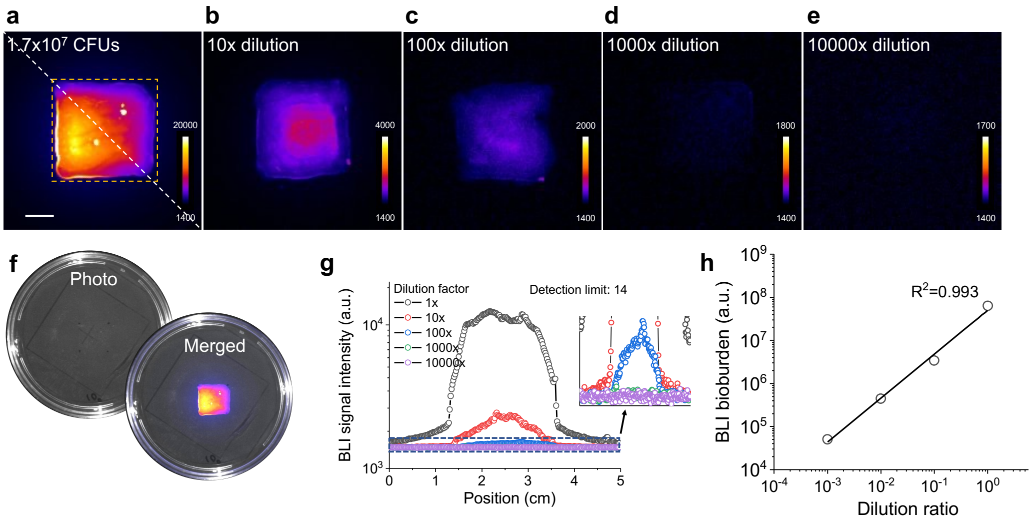


**Figure S7.** Measurement to confirm the linearity between bioluminescence (BLI) signal and bacterial load. (a-e) BLI images of *P. aeruginosa* PAO1 *lux* cells inoculated on agar plates with different dilution factors thus different bacterial load. The starting inoculation load in (a) was 1.7×10$^7$ CFUs, which was uniformly smeared within an area of 15×15 mm$^2$ (yellow dashed frame in (a)) simulating the bacterial inoculation on a rat wound. Scale bar, 5 mm. The BLI reached its detection limit at a dilution factor of 10000× in (e). (f) Photo of the *in vitro* agar plate in (a) and a merger with its BLI image. (g) BLI signal plot for (a-e) along a diagonal axis (white dashed line in (a)). Inset, a zoom-in plot. The detection limit was 14 CFUs for *P. aeruginosa* PAO1 *lux* under the imaging setting. (h) The integrated BLI signal intensity or total BLI bioburden for (a-e), showing a linear relationship with the bacterial load.

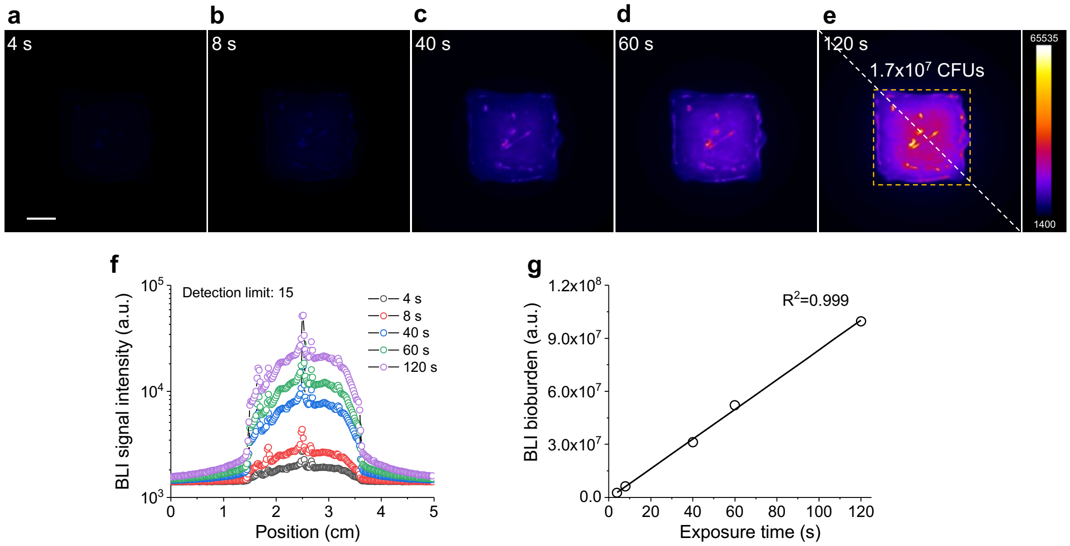


**Figure S8.** Measurement to confirm the linear relationship between bioluminescence (BLI) signal and camera integration time for *P. aeruginosa* PAO1 *lux* cells. (a-e) BLI images of bacterial cells inoculated on an agar plate under different integration times. The inoculation load was 1.7×10$^7$ CFUs, which was uniformly smeared within an area of 15×15 mm$^2$ (yellow dashed frame in (e)) simulating the bacterial inoculation on a rat wound. Scale bar, 5 mm. (f) BLI signal plot for (a-e) along a diagonal axis (white dashed line in (e)). (g) The integrated BLI signal intensity or total BLI bioburden for (a-e) over the camera exposure time or integration time.

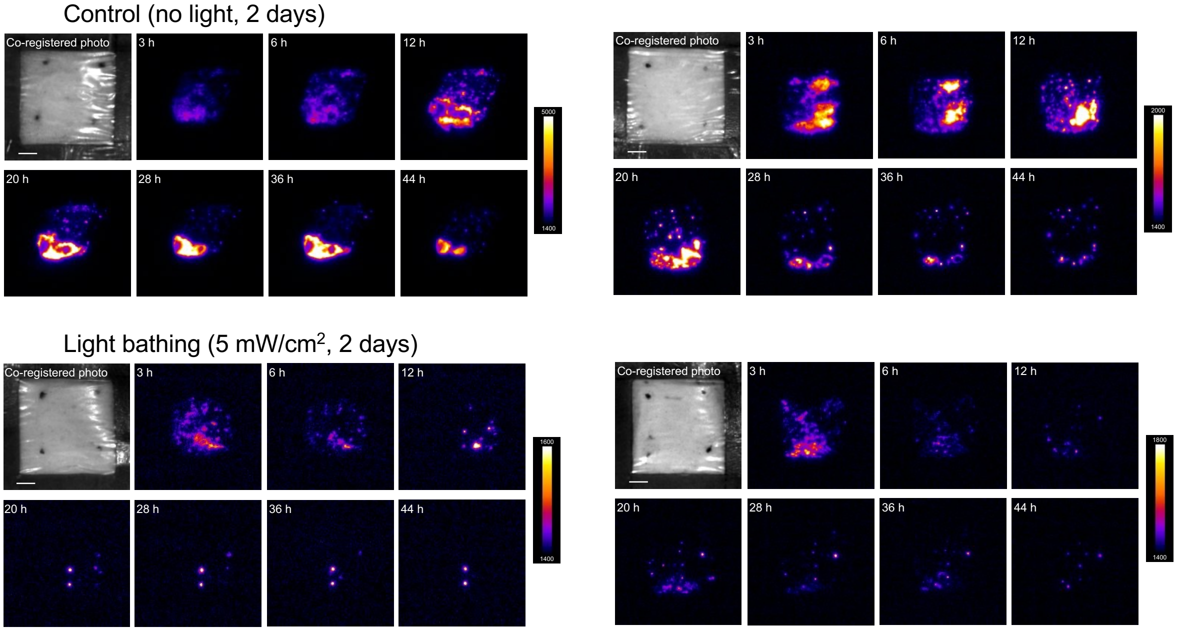


**Figure S9.** Time-lapse bioluminescence imaging (BLI) data of untreated (top) and treated (bottom) MRSA USA300 *lux*-infected abrasion wounds on two more rats. (Left) Second set of independent replicates. (Right) Third set of independent replicates. The first image in each set showed the photo of the infected wound. Scale bar, 5 mm.

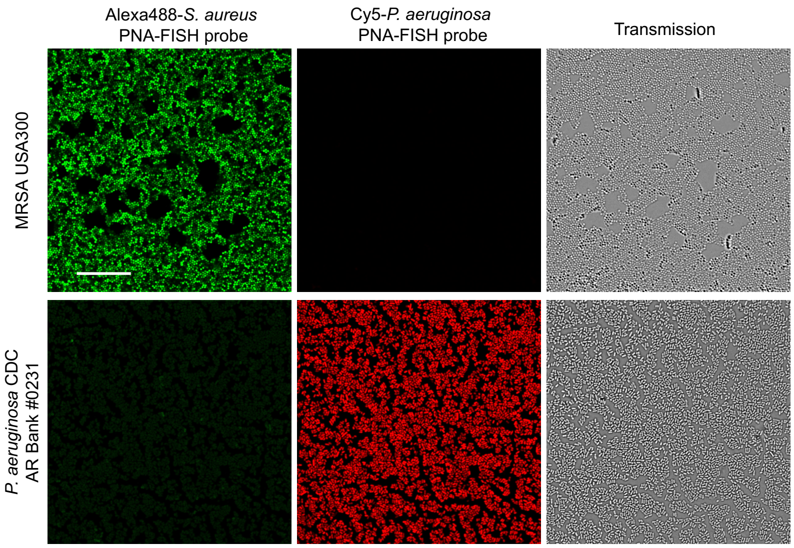


**Figure S10.** PNA-FISH imaging of *S. aureus* (MRSA USA300) and *P. aeruginosa* (CDC AR Bank #0231) *in vitro* for specificity validation. No crosstalk was confirmed between the two bacterial species-specific probes. Scale bar, 30 µm.

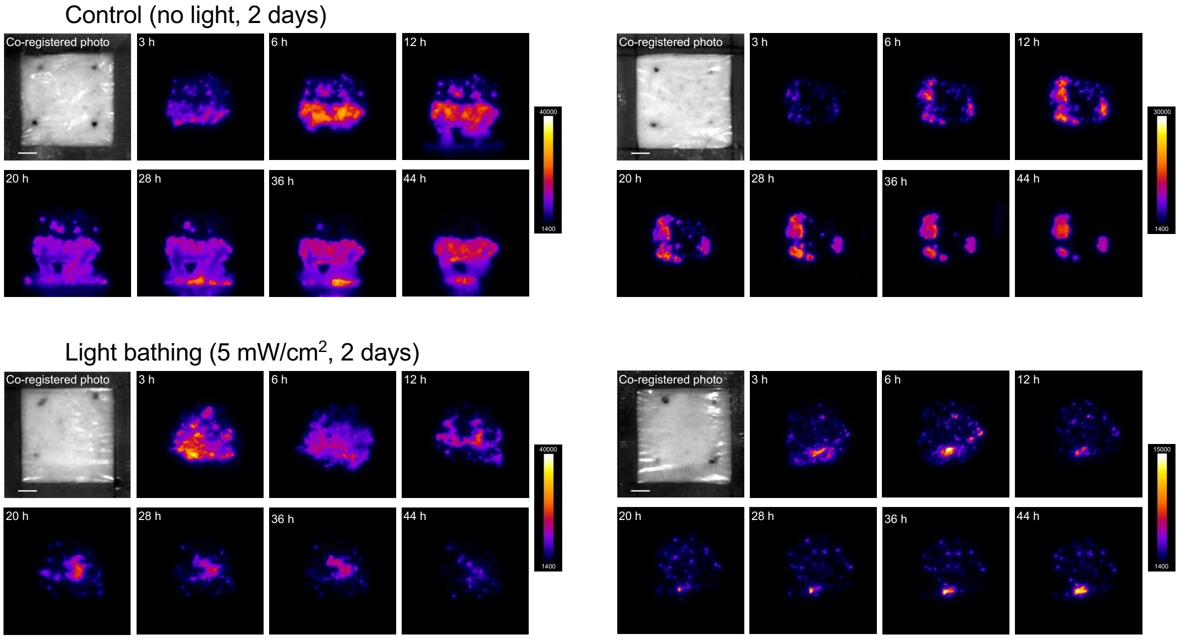


**Figure S11.** Time-lapse bioluminescence imaging (BLI) data of untreated (top) and treated (bottom) *P. aeruginosa* PAO1 *lux*-infected abrasion wounds on two more rats. (Left) Second set of independent replicates. (Right) Third set of independent replicates. The first image in each set showed the photo of the infected wound. Scale bar, 5 mm.

### Descriptions of Supplementary Videos

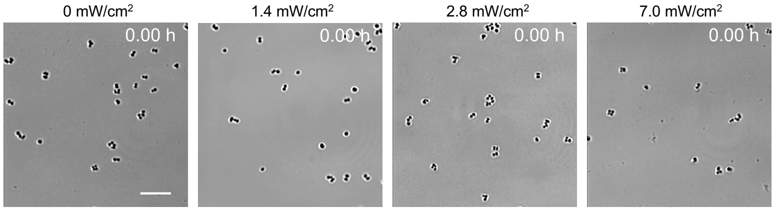


**Video S1.** Time-lapse confocal images of MRSA USA300 cells *in vitro* with concurrent, continuous blue light illumination at different irradiances. Scale bar, 20 μm.

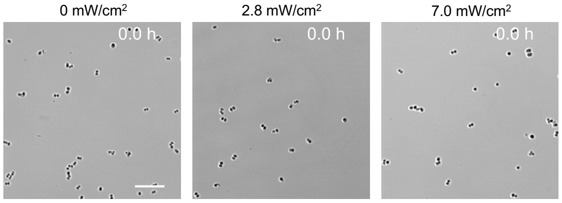


**Video S2.** Time-lapse confocal images of MRSA USA300 cells *in vitro* supplemented with 16 mM DMTU and further with concurrent, continuous blue light illumination at different irradiances. Scale bar, 20 μm.

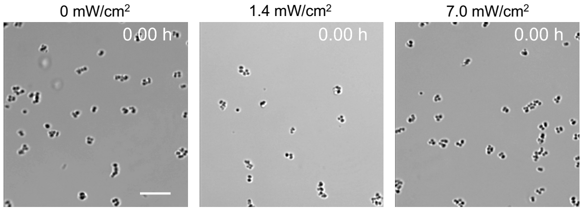


**Video S3.** Time-lapse confocal images of catalase-deficient mutant *S. aureus* Δ*katA* cells *in vitro* with concurrent, continuous blue light illumination at different irradiances. Scale bar, 20 μm.

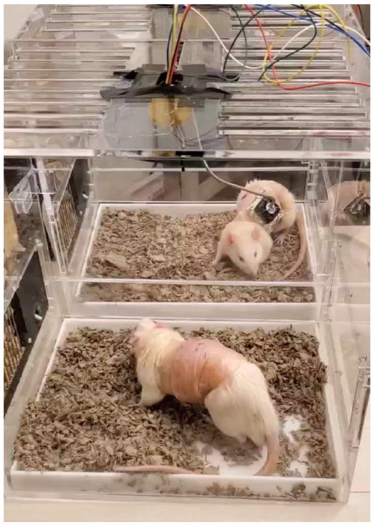

**Video S4.** Video of rat models used in the study with and without a light-delivering wound patch.
